# Hypoxia in cancer chemo- and immunotherapy: foe or friend?

**DOI:** 10.1101/629907

**Authors:** Noemi Vitos, Shannon Chen, Shreya Mathur, Ibrahim Chamseddine, Katarzyna A. Rejniak

## Abstract

Hypoxia, a low level of oxygen in the tissue, is a feature of most solid tumors. It arises due to an imbalance between the oxygen supply from the abnormal vasculature and oxygen demand by the large number of tumor and stromal cells. Hypoxia has been implicated in the development of aggressive tumors and tumor resistance to various therapies. This makes hypoxia a negative marker of patients’ survival. However, recent advances in designing new hypoxia-activated pro-drugs and adoptive T cell therapies provide an opportunity for exploiting hypoxia in order to improve cancer treatment. We used novel mathematical models of micro-pharmacology and computational optimization techniques for determining the most effective treatment protocols that take advantage of heterogeneous and dynamically changing oxygenation in in vivo tumors. These models were applied to design schedules for a combination of three therapeutic compounds in pancreatic cancers and determine the most effective adoptive T cell therapy protocols in melanomas.

## I. Introduction

Tumor tissues are characterized by low oxygenation (often 5-10 times lower than in normal tissues), which emerges due to limited oxygen perfusion and/or diffusion. As a result, the tumor tissue can contain multiple hypoxic regions, each with a possibly different level of O_2_. Hypoxia is associated with aggressive tumor behavior, metastatic spread capabilities, and resistance to chemo-, immuno-, and radiation therapies [1]. To overcome the therapeutic barriers created by hypoxia, novel chemotherapeutic compounds are being developed that become chemically active only in areas of low oxygen levels and remain non-lethal in the well-oxygenated regions. These treatments are known as hypoxia-activated pro-drugs (HAPs) [2]. Another approach, known as adoptive cell therapy (ACT), uses patients’ immune cells for in vitro enhancement and transfer back to the patient’s blood circulation to improve the anti-tumor immune responses [3]. While both these types of treatment are currently in clinical trials for multiple solid tumors, there is still a need for methods to improve their efficiency. Here, we propose using a mathematical modeling framework of micro-pharmacology to design the optimal protocols for HAPs and ACT.

## II. Designing multi-chemotherapy schedules for exploiting oxygenation levels in pancreatic tumors

Hypoxia presents a barrier to effective chemotherapy, since the hypoxic tumor cells remain quiescent and stop proliferating. Thus, drugs that target rapidly proliferating cells have become ineffective. HAPs can provide a way for overcoming this hurdle. However, the clinical efficacy of these drugs could be improved by acutely exacerbating tumor hypoxia to increase the area and time window of HAP activity. Here, we consider two previously tested compounds, namely a vasodilator that temporarily decreases blood (and oxygen) flow to the tumor vasculature [4] and a metabolic sensitizer that increases oxygen consumption by the cells exposed to high concentrations of it [5]. Using the micropharmacology model, *microPKPD* (Figure 1A), we systematically tested the various combination schedules of HAP with a vasodilator alone, sensitizer alone, and both compounds together for identifying the most effective protocols.

**Figure 1.**
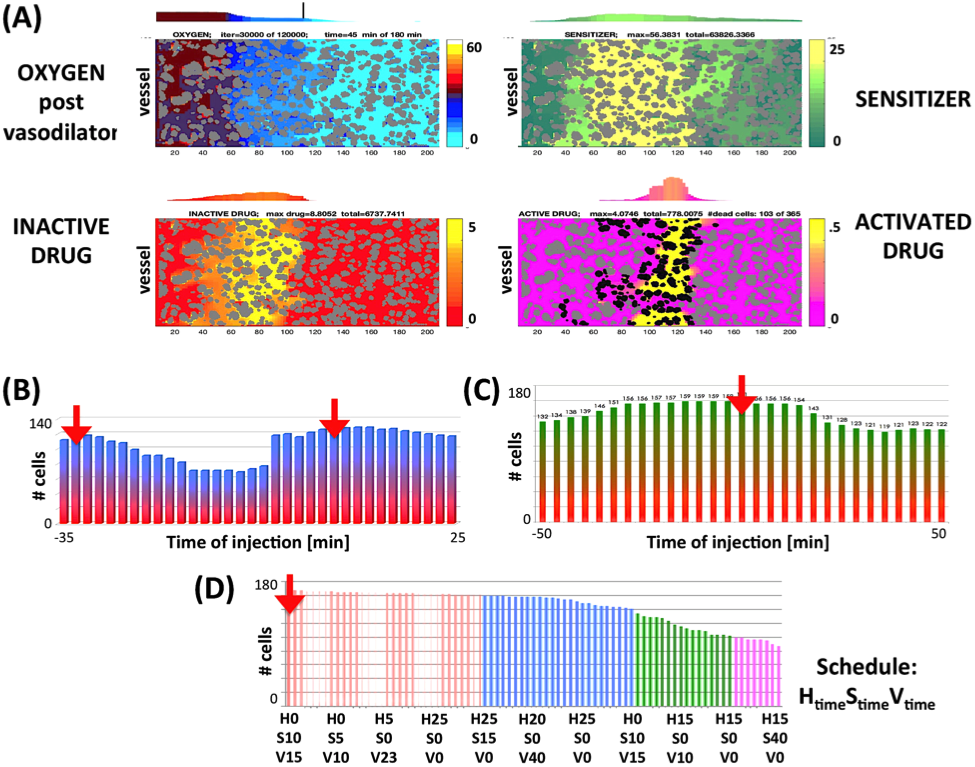
(A) The *microPKPD* model for testing the efectiveness of drug combinations: oxygen after vasodilator injection (top-left), sensitizer (top-right), inactive pro-drug (bottom-left), and activated drug (bottom-right). Each colobar shows the corresponding concentration level (low-bottom color & high-top color). (B-C) Two-tretment combination schedules: HAP+vasodilator (red-blue, B) and HAP+sensitizer (red-green, C), colors correspond to color-scheme in panel (A). (D) Three compound treatment schedules ordered from the most to least effective. Red arrows show the optimal schedules.

Our simulated results of two-drug therapies indicated that the best outcomes were achieved when the vasodilator was injected either shortly after HAP or sufficiently long before HAP to avoid a reduction in HAP influx from the dilated vasculature (two peaks in Figure 1B), and when the sensitizer was administered prior to HAP (plateau in Figure 1C). Interestingly, for three-drug combinations, the injection of HAP first followed by the sensitizer and vasodilator resulted in doubling the number of dead tumor cells when compared with injection of HAP alone, while a 25% dead cell increase occurred compared with two-drug combinations.

## III. Designing adoptive T cell therapy protocols for exploiting oxygen heterogeneity in melanomas

Adoptive T cell therapy (ACT) promises to be an effective personalized treatment, since it involves the transfer of host immune cells after they are enhanced in vitro. This process requires growing large numbers of antitumor lymphocytes and selecting those with high-avidity recognition of the tumor. The selected tumor-infiltrating lymphocytes (TILs) are then injected back into the patients. However, there is room for improving the clinical efficacy of this approach. We used the micro-pharmacology model, *microPKPD* (Figure 2A), for testing ways of increasing tumor tissue penetration by the TILs and the interferon gamma (IFNγ)-dependent interactions between TILs and tumor cells [6]. Our laboratory experiments showed that TILs’ viability is not affected by low levels of oxygen, but others have reported that TILs activated under hypoxia produce higher amounts of IFNγ. Thus, we tested several ways of increasing IFNγ distribution in the tumor tissue.

**Figure 2.**
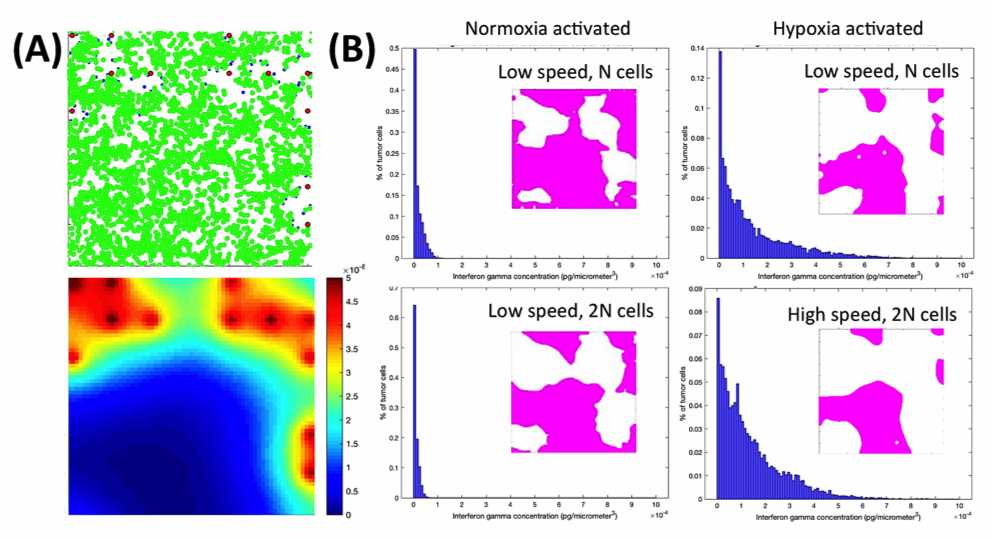
(A) The *microPKPD* model for testing the effectiveness of ACT-TIL: shown are (top) tissue topology with tumor cells (green), vessels (red) and infiltratinf T cells (blue) and (bottom) oxygen distribution with the colorbar indicating hypoxia (blue), normoxoa (cyan to orange) and intravascular O_2_ level (red). (B) A histogram showing IFNγ distribution around the tumor cells (blue) and within the tissue (pink) for low- and highspeed T cells with regular and doubled numbers of T cells that were either normoxia- or hypoxia-activated.

Our simulations showed that doubling the number of TILs did not increase the spatial distribution of IFNγ in the tumor tissue. In contrast, the hypoxia-activated TILs resulted in significantly increased levels of IFNγ both in the tissue, and specifically around the tumor cells. Moreover, when the in silico T cells were migrating with double speed, the IFNγ concentration around tumor cells was elevated.

## IV. Discussion

We presented here two applications of the *microPKPD* model to test how heterogeneity in tumor tissue can be exploited for improving anti-cancer therapies. While tumor hypoxia is typically associated with a poor treatment prognosis, the recently developed treatments take advantage of various levels of oxygen. Our simulations examined how to improve the efficacy of these treatments by designing optimal schedules of hypoxia-activated drug combinations and predicting optimal activation protocols for the TIL adoptive cell therapy (ACT-TIL).

## V. Mathematical Methods

The computational micro-pharmacology model, *microPKPD* considers the topology of the tumor tissue and pharmaco-kinetic/pharmaco-dynamic (PK/PD) properties of drugs and chemokines secreted by the T cells. The model couples reaction-advection-diffusion equations with fluid dynamics equations and particle-spring model equations. The drug and chemokine kinetics (Eqs.1,2) are described by reaction-advection-diffusion equations for the HAP model and reaction-diffusion (no advection part) for the ACT-TIL model. The advection part is calculated by the regularized Stokeslets method (Eqs.3,4). A movement of an individual T cell is governed by a combination of forces (Eq. 5): a random migration drag force (Eq.6) and a passive repulsive force that prevents the cell from overlapping with other T cells and tumor cells (Eqs.7). Model parameters are listed in Table 1. More details are given in [7].

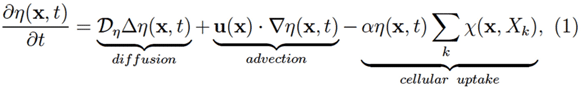

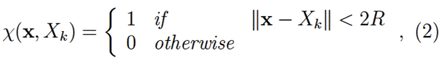

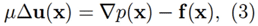

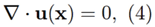

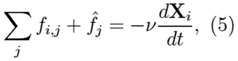

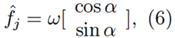

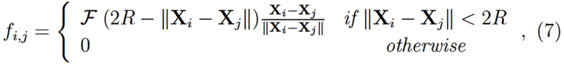

**Table 1.** Parameters from the *microPKPD* model of hypoxia-activated prodrugs (top, HAP) and adoptive cell therapy with tumor infiltrating lymphocytes (bottom, ACT-TIL).

| parameter | value |
| --- | --- |
| **HAP model** |  |
| oxygen diffusion | $\mathcal{D}_o = 10^{-6} \text{ cm}^2/\text{s}$ |
| inactive drug diffusion | $\mathcal{D}_{\eta_i} = 5 \times 10^{-7} \text{ cm}^2/\text{s}$ |
| active drug diffusion | $\mathcal{D}_{\eta_a} = 2 \times \mathcal{D}_{\eta_i}$ |
| sensitizer diffusion | $\mathcal{D}_\gamma = \mathcal{D}_o$ |
| interstitial fluid velocity | $\mathbf{u} = 2 \mu\text{m}/\text{s}$ |
| active drug uptake | $\alpha_{\eta_a} = 0.5 \text{ pg}/\text{s}$ |
| tumor cell radius | $\mathcal{R} = 10 \mu\text{m}$ |
| **ACT-TIL model** |  |
| oxygen diffusion | $\mathcal{D}_o = 10^{-6} \text{ cm}^2/\text{s}$ |
| cytokine diffusion | $\mathcal{D}_c = 10^{-8} \text{ cm}^2/\text{s}$ |
| T cell radius | $\mathcal{R} = 5 \mu\text{m}$ |
| T cell speed | $\omega = 8 \mu\text{m}/\text{s}$ |
| repulsive spring stiffness | $\mathcal{F} = 50 \mu\text{g}/\mu\text{m} \cdot \text{s}^2$ |
| medium viscosity | $\nu = 250 \mu\text{g}/\mu\text{m} \cdot \text{s}$ |

